# Individual reflectance of solar radiation confers a thermoregulatory benefit to dimorphic males bees (*Centris pallida*) using distinct microclimates

**DOI:** 10.1101/2022.06.28.497958

**Authors:** Meghan Barrett, Sean O’Donnell

## Abstract

Incoming solar radiation (wavelengths 290 – 2500 nm) significantly affects an organism’s thermal balance via radiative heat gain. Species adapted to different environments can differ in solar reflectance profiles. We hypothesized that conspecific individuals using thermally distinct microhabitats to engage in fitness-relevant behaviors would show intraspecific differences in reflectance: we predicted individuals that use hot microclimates (where radiative heat gain represents a greater thermoregulatory challenge) would be more reflective across the entire solar spectrum than those using cooler microclimates. Differences in near-infrared (NIR) reflectance (700 - 2500 nm) are strongly indicative of thermoregulatory adaptation as, unlike differences in visible reflectance (400 - 700 nm), they are not perceived by ecological or social partners. We tested these predictions in male *Centris pallida* (Hymenoptera: Apidae) bees from the Sonoran Desert. Male *C. pallida* use alternative reproductive tactics that are associated with distinct microclimates: Large-morph males, with paler visible coloration, behave in an extremely hot microclimate close to the ground, while small-morph males, with a dark brown dorsal coloration, frequently use cooler microclimates above the ground near vegetation. We found that large-morph males had higher reflectance of solar radiation (UV through NIR) resulting in lower solar absorption coefficients. This thermoregulatory adaptation was specific to the dorsal surface, and produced by differences in hair, not cuticle, characteristics. Our results showed that intraspecific variation in behavior, particular in relation to microclimate use, can generate unique thermal adaptations that changes the reflectance of shortwave radiation among individuals within the same population.

## Introduction

Solar (shortwave) radiation plays a large role in an organism’s thermoregulation via radiative heat gain, driving adaptive patterns of body coloration and reflectance [1–2]. Organisms may absorb solar energy to heat up, or transmit/reflect solar energy to avoid additional heat gain [3–4]. Environmental conditions are particularly important for ectotherms, which rely heavily on managing the influence of abiotic factors like shortwave radiation to maintain non-lethal body temperatures [5]. Differences in reflectance across the entire solar ultraviolet (UV) to near infrared (NIR) spectrum can thus have significant thermoregulatory consequences [6–7]; for example, the reflective hairs of Saharan silver ants reduce their body temperatures by as much as 7-10 °C [8]. The thermal effects of variation in solar radiation affect population and species variation in color and reflectance across landscapes [2,4,9–12].

Differences in ectotherms’ visible coloration (henceforth, VIS reflectance, wavelengths 400 - 700 nm) can serve as a morphological mechanism for thermal adaptation (e.g. Thermal Melanism Hypothesis or Bogert’s Rule [13–16]). External coloration affects operative temperatures both across and within lizard species [17–18]. However, climate is a poor predictor of visible color in butterflies [2, but see 11], and ant assemblages demonstrate conflicting macrophysiological coloration patterns [19–20]. These conflicting patterns may result from the fact that VIS (and UV; 290-399 nm) reflectance is under selection by factors not related to thermoregulation, including social/sexual signaling and anti-predator adaptations, which may conflict with thermal selective effects [1,16]. By contrast, differences in external reflectance of NIR (701 – 2500 nm) radiation are not confounded by ecological and social partners, and are thus generally only under selection for thermoregulatory reasons [1]. NIR reflectance has larger impacts on heating rates than UV and VIS reflectance [7], and is more consistently associated with climate variation [2,4,8,11].

Research on adaptive coloration and reflectance has focused on population or species differences [2,11,19–20]. However, there can be significant differences in body temperature associated with radiative heat gain in different microhabitats (e.g., sunny vs. shaded sites [21]). Within populations, individuals may vary consistently in which microhabitats they use based on their behaviors, with significant thermoregulatory consequences [21–25]. These individual differences are overlooked by population- and species-level studies, which treat conspecific individuals as ecologically equivalent despite the fact that individual variation can affect ecological and evolutionary dynamics and serves as a major target of natural selection [26]. Studies that measure within-population differences can yield new insights into the selective pressures that act on individuals to generate evolutionary adaptations to fine-scale thermal variation.

*Centris pallida* bees live in the Sonoran Desert and experience extremely high temperatures during their mating aggregations [27]. Males are dimorphic in behavior and morphology: large-morph males patrol near the ground, fully exposed to the sun, and attempt to locate females in underground nests using scent. Large-morph males then dig out and fight over females, which can last 19 minutes or more [28]. Small-morph males occasionally participate in this strategy, but more typically hover 1 m or more above the ground near vegetation and chase after flying females they locate visually [29–31]. Air temperature in the typical large male microclimate is nearly > 8 °C hotter than the typical small male microclimate by the time mating behaviors cease, around 1000 - 1100 hr [32].

Large-morph males are paler in visible coloration than small-morph males and females (Fig 1); however, higher VIS reflectance may be driven in part by the need to camouflage themselves against the desert soil to reduce predation events by birds and lizards [33]. If the higher reflectance of large-morph males extends into the NIR, it likely represents a thermoregulatory adaptation to reduce their radiative heat gain in the hotter microclimate. The darker coloration of small-morph males may also be thermally adaptive, allowing them to maximize performance during cooler early-morning periods when males are observed to bask on bushes prior to foraging/mating around 0700 (Barrett, pers. comm.). Radiative heat gain is critical at low air temperatures (particularly for flying insects that lose heat via convective cooling [24]), and plays a large role in allowing ectotherms to reach active temperatures at cool air temperatures [34]. Females may also be darker in coloration (like small-morph males) in order to increase solar radiative heat gain and maximize foraging performance in the cool, early-morning period.

**Fig 1.**
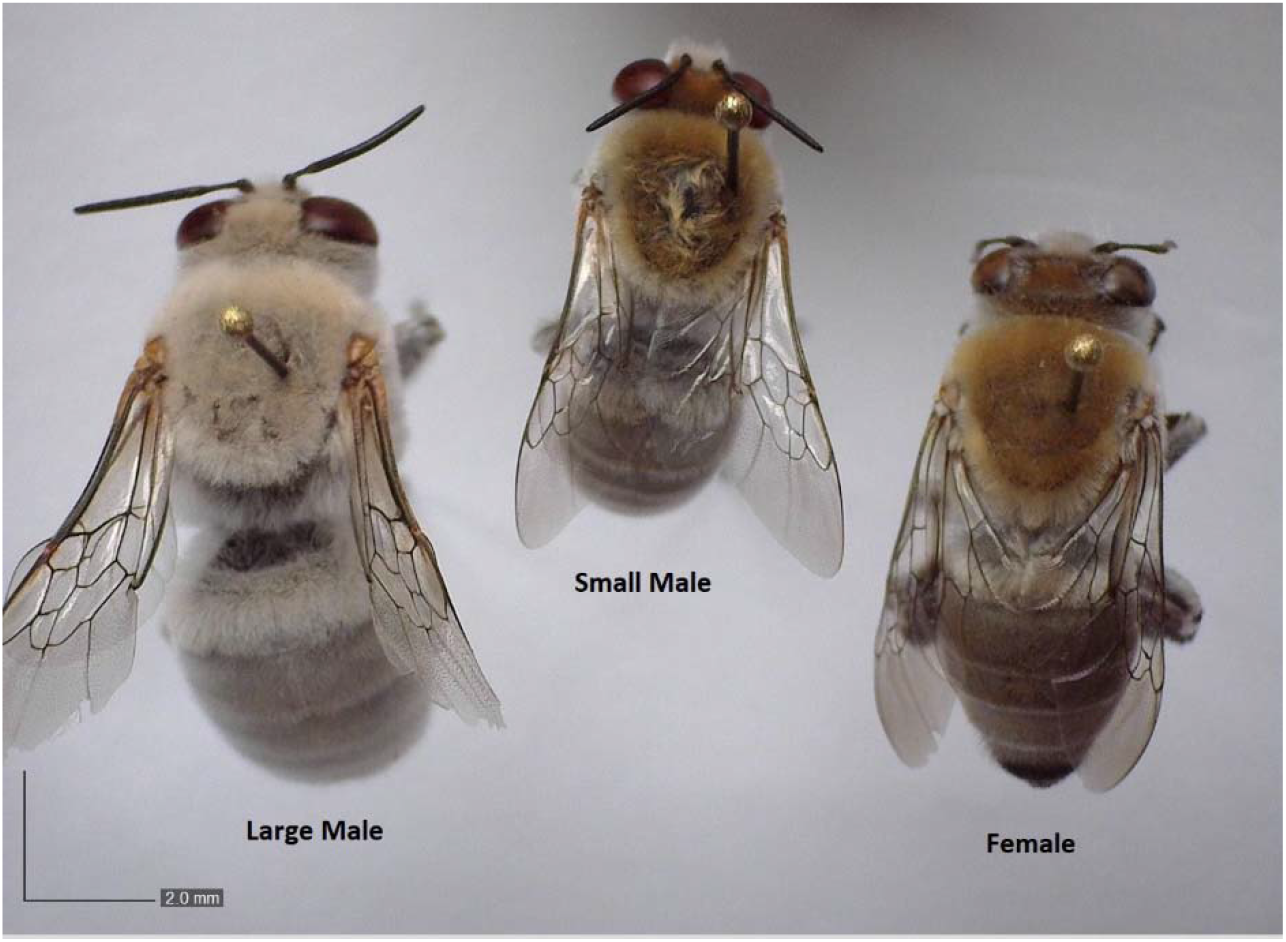
Visible coloration and size dimorphism in *C. pallida*. Large-morph *C. pallida* male (left) has a larger body size and a pale-grey coloration across the head, thorax, and abdomen compared to the small-morph *C. pallida* male (middle), which has a brown head and thorax and dark grey abdomen, most similar to the female of generally intermediate size (right). Close-up photos of thorax and abdomen coloration in S1 Fig.

We hypothesized that microclimate usage drives morph-specific differences in UV-NIR reflectance. We predicted that large-morph males would have higher reflectance of solar radiation compared to small-morph males, as a thermoregulatory adaptation to reduce or increase shortwave radiative heat gain in their hotter or cooler microclimates, respectively. Because microhabitat differences are strongest in incident radiation from above, we predicted morph reflectance differences on the dorsal, but not ventral, body surfaces (see [8]). To better understand mechanisms of morph-specific reflectance we tested whether differences in reflectance were related to hair (branched setae) or cuticle characteristics; we measured solar reflectance of the dorsal surface of the thorax with, and without, hairs present. We predicted that hair density would correlate with higher reflectance across body regions, or morph/sex, and calculated hair density on the thorax and abdomen across morphs and females. Finally, we predicted that females would have similar reflectance profiles to small-morph males on the dorsal surface, and would not differ from males in ventral surface reflectance.

## Materials and methods

### Specimen Collection

We classified males as the ‘large’ male morph if we found them patrolling, digging, or fighting and they had the grey/white coloration (Fig 1, S1 Fig) and leg morphology (thick, bulging femurs on the hindmost pair of legs) distinctive to the large, behaviorally-inflexible morph [28–39,35]. We classified them as ‘small’ male morphs if we collected them hovering near vegetation (large-morph males never engage in hovering behavior [28]). We collected *C. pallida* males and females (n = 10/morph or sex) in late April and early May of 2018, or in late April and early May of 2019, either within 10 km of N33.464, W111.632 or N32.223, W11.008. Permits were obtained through the Desert Laboratory on Tumamoc Hill, or bees were collected non-commercially on public lands and no permits were required. We transported bees in a cooler on ice to a lab where we weighed them on an analytical balance (Metler Toledo AB54-S) to the nearest 0.1 mg.

### Morphological Measurements

We used digital calipers (Husky, 6 inch - 3 mode model) to measure the body parts of the male and female bees used for reflectance spectrophotometry. We assumed the thorax and head to be a single cylinder for surface area calculations, with a diameter was equal to the widest point of the thorax and height equal to the length of the bee from the top of the head to the back of the thorax. We assumed the abdomen was a cylinder with a diameter equal to the width of the second tergite, and height equal to the length of the abdomen. For surface area calculations, we subtracted one flat, circular side (e.g., base) of both the thorax-head, and abdomen, cylinders, in order to account for the surface where the abdomen and thorax meet.

### Reflectance Spectrophotometry

We stored males of each morph and females dry, and away from light, following field collection. We captured total diffuse and spectral ultraviolet to near-infrared (290 - 2500 nm) percent reflectance (R) of the external surface using a Cary 5000 spectrophotometer equipped with an Agilent integrating sphere, deuterium arc (<350 nm) and tungsten-halogen (>350 nm) lamps, and R928 photomultiplier and PbS IR detectors. We limited measurements to a consistent diameter for all specimen using a custom aluminum aperture (2.18 mm diameter, 0.66 mm thickness), which we also used during the zero and baseline (Labsphere certified reflectance standard, Spectralon) measurements. We set integration time to 0.2 seconds, with data intervals of 1 nm. Baseline measurements varied across set up days, possibly due to minor contamination on the Spectralon surface, so we used the following formula to standardize measurements (R_standardized,n_) across different set-ups:

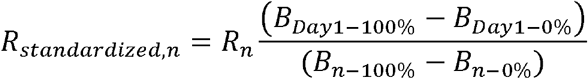

Where R_n_ is the measured percent reflectance on Day n, B_100%_ is the Spectralon baseline (on Day 1 through Day n) and B_0%_ is the zero baseline (on Day 1 through Day n).

We measured R on the dorsal surface of the thorax (unshaved, and shaved of all hairs using a razor blade and the tips of #5 forceps), the dorsal surface of the abdomen on the first two terga, and the ventral surface of the abdomen on terga 2-3, for each specimen (n = 10 individuals of each morph/females for each area). We mounted specimen so that the thorax or abdomen was flat against the aperture, with measurements normal to the surface. Measurement error occurred between 799 and 800 nm due to the detector switching from the R928 to PbS IR (an artefact of sample orientation changing in relation to the new detector). To account for this, we took the difference in R between 799 and 800 for each individual and then added half of this value to all measurements ≤ 799 and subtracted it from all values ≥ 800. As *C. pallida* can lose hair over their lifetime, we only used individuals collected when the aggregation had just started, with low wing wear and no visible hair thinning on the thorax.

We assumed transmission through the bee was 0 (a standard assumption for insect bodies in the NIR [6,8]), and calculated average absorption (*a_n_*) for a morph or sex at a wavelength ‘n’, using the following equation:

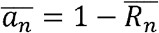

where 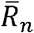 is the average fraction of reflectance of all ten specimen at wavelength n.

### Solar Radiation Calculations

Thermal flux due to solar radiation can be defined by summing direct beam (b), diffuse sky (d), and reflected (from substrates; r) radiation:

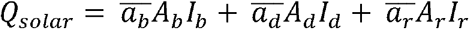

where *ā* is the absorption coefficient of the bee (the fraction of beam, diffuse, or reflected radiation absorbed by the bee); A_b_, A_d_, and A_r_ are the surface area of the bee exposed to beam, diffuse, or reflected radiation; and *Ib, Id*, and *I_r_* is the total irradiance of the beam, diffuse, or reflected source. Here, we assumed A_b_ = 0.25 A_s_ (total surface area), and A_d_ and A_r_ = 0.5 A_s_ (standard assumption for bees, and cylindrical objects, as in [36]). For A_b_ and Ad, we used the total thorax and abdomen surface areas separately for As, in order to take advantage of the two distinct absorption coefficients we calculated for each region dorsally; for A_r_, we combined the two regions into a total surface area. Because only male morphs consistently differ in surface area, we did not perform thermal flux calculations for females; we did, however, calculate their absorption coefficients as described below.

We obtained the direct normal irradiance spectra in order to calculate 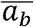 and 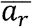 from the National Renewable Energy Laboratory (ASTM G173-03 Reference Air Mass 1.5 Spectra Derived from SMARTS v. 2.9.2 [37]). We calculated the direct beam absorption coefficient (a_b_) for each individual on the dorsal surface of the abdomen and the thorax, as well as the shaved cuticle, between 290 and 2500 nm λ_min_ and λ_max_) as:

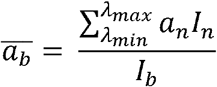

Where *I*_n_ is the intensity of solar radiation at wavelength n and *I*_b_ is the total direct beam irradiance between the min and max wavelengths.

We obtained the spectrum of radiation reflected from a light soil substrate (RS^substrate^), at each wavelength (n), from the USGS Spectral Library (sample ID: splib07a record = 13249 [38]), in order to calculate the intensity of reflected radiation, *I*_r_, based on the intensity of direct beam solar radiation (*I*_b_) at each wavelength (n).

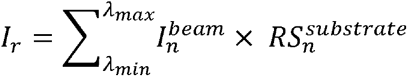

We then calculated the average absorption coefficient 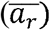 of reflected radiation, using each bee’s ventral abdominal absorption (a_rn_) coefficient at each wavelength (n):

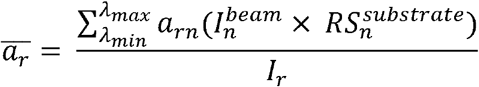

We also calculated a second measure of *I_r_* using an albedo measurement of light sand from the Sonoran Desert (*I*_r_ = 0.245*I*_b_ [39]), to compare with the light soil substrate spectrum obtained from the USGS spectral library. Unlike the USGS sample, this summed-reflectance value could not account for wavelength specific effects – but this value had the advantage of being from the location of interest.

We assumed 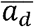 to be equal to 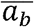 (as in [36]). We assumed *I*_d_ was a 100 W/m^2^ [40]. We estimated direct beam radiation on April 20^th^, 2018 at 1000 hr using the following formulas from the solar radiation geometry literature [41–43]:

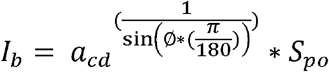

With the following assumptions: S_po_=1361 W/m^2^, a_cd_ =0.83 (clear day), and atmospheric pressure at the study site = standard air pressure (101.3 x 10^3^ kPa). Altitude angle (ø) was calculated using the following equations, using the Julian date (J = 110), Latitude (32.2°, 0.562 rad [λ]), time (t = 1000), and solar noon time (t_0_ = 1200):

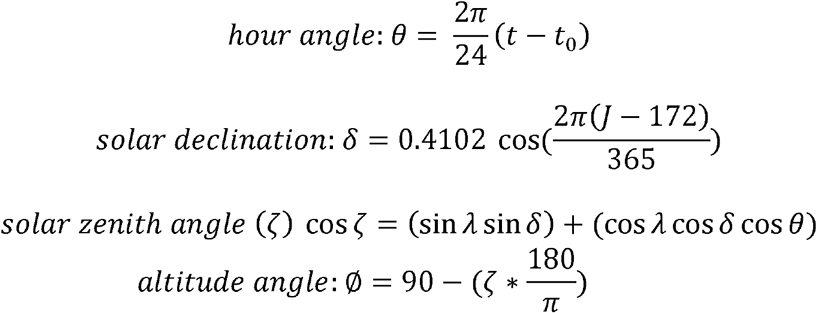

Finally, We divided Q_solar_ by body mass (g) to determine solar radiative heat flux per unit of body mass.

### Scanning Electron Microscopy

We shaved small patches of the second tergite of the abdomens of the specimen used for reflectance spectrophotometry (thoraxes were already shaved). We then mounted thoraxes and abdomens on metal stubs with electrically conductive tape and silver paint, before sputter-coating samples (Pt/Pd target, 80/20; Cressington 208 Hr Sputter Coater) for 40 s at 40 mA (approximately 8-10 nm deposition). We took three photos of the cuticular surface in three different places on the abdomen/thorax at 100X or 200X magnification using a Zeiss Supra 50VP (EHT set to 5 kV, high vacuum mode, SE2 detector). We used ImageJ to count the number of pores found at the base of the hairs within a standardized, circular region of 0.1 mm^2^ area (centered on a pore) on each of the three photos for each individual, and averaged these three numbers to obtain hair density in terms of the number of hairs/mm^2^. On the abdomen, both unbranched and branched setae can be found, however the pores that at the base of these two setae types are quite distinct (Fig 2); we counted only the pore type that led to the branched setae, which was more common (in photo 2A, for instance, only 2.9% of pores are for unbranched setae).

**Fig 2.**
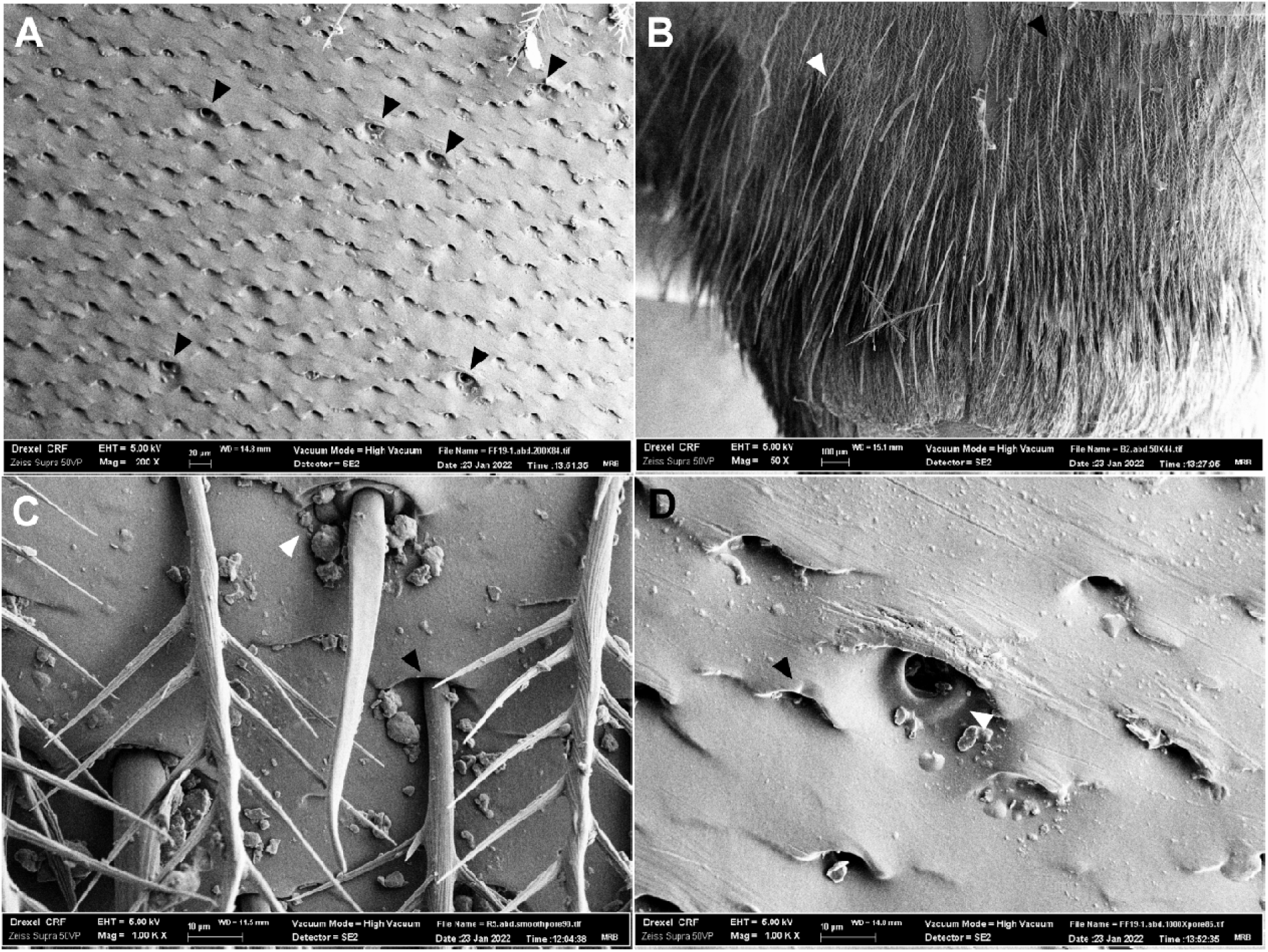
Difference between pores of branched (hairs) and unbranched setae on the abdomen. A) Female abdominal cuticle, dorsal surface, with black arrowheads pointing to the locations of unbranched setae pores (200X). This photo is representative of those used for quantification of hair density (taken on Terga 2). B) Terga 5 & 6 of a female *C. pallida* showing many unbranched setae (white arrowhead) against a background of generally shorter branched setae (black arrowhead; 50X). Unbranched setae appeared longer and more common on terminal terga. C) Female abdominal cuticle, showing difference in pores with branched (black arrowhead) and unbranched (white arrowhead) setae still attached on Terga 2 (1000X). D) Close-up of (A), showing difference in branched (black arrowhead) and unbranched (white arrowhead) setae pores (1000X).

### Analyses

All data are available on Dryad (DOI: https://doi.org/10.5061/dryad.73n5tb31q). We used GraphPad Prism 9.1.2 [44] and and R v. 4.1.3 [45] to analyze spectrophotometry, morphology, and hair density data. We used one-way ANOVAs with Tukey’s MCT to compare body mass, and thorax and head or abdominal surface area across large/small males and females. We used a one-way ANOVA followed by Tukey’s MCT for normally distributed data, or Kruskal-Wallis test with Dunn’s MCT for data that were not normally distributed, to assess variation in hair density on the abdomen and thorax. We calculated mean percent reflectance in the UV (290-399), VIS (400-700) and NIR (split into “close NIR” [cNIR, 701 – 1400] and “far NIR” [fNIR, 1401 – 2500]) for each individual, and used a two-way RM ANOVA (with a Geisser-Greenhouse correction due to lack of sphericity in wavelength) to assess the affect of morph (e.g. large-morph male, small-morph male, and female), wavelength region (UV, VIS, cNIR, and fNIR), and individual differences in mean reflectance. When we found significant p-values for ‘morph’, we used a Tukey’s MCT to assess variation across categorical morph assignments within each wavelength region (and adjusted p-values for multiple comparisons). To test for differences in mean reflectance in the UV, VIS, cNIR, and fNIR between the shaved and unshaved dorsal thorax surface, or the dorsal and ventral side of the abdomen, of large and small males, we used two-way RM ANOVAs with Geisser-Greenhouse corrections. When we found significant p-values for the effect of ‘shaving’ (thorax) or ‘side’ (abdomen), we used a Sidak’s MCT to assess variation associated with that variable within each wavelength region (and adjusted p-values for multiple comparisons). To test for a specific thermal adaptation across morphs/sexes in the NIR on the dorsal surface of the thorax and abdomen, while controlling for correlations with VIS reflectance, we tested the morph/sex and mean VIS reflectance for each individual as fixed effect predictors of NIR reflectance in a linear model, using repeated single-term deletions based on AIC comparisons followed by Type 1 ANOVA p-value comparisons to determine the simplest, best-fit model. We used one-way ANOVAs followed by Tukey’s MCTs were used to assess differences in absorption coefficients between morphs and females.

## Results

### Morphology & Hair Density

Large and small morph males, and females differed in their wet body masses (Table 1; One-way ANOVA: F = 42.80, df = 32, p < 0.0001). Large-morph males had larger abdomen, and thorax + head, surface areas compared to both small-morph males and females (Table 1; thorax and head SA: F = 53.70, df = 32, p < 0.0001; abdomen SA: F = 40.25, df = 32, p < 0.0001); small-morph males had slightly smaller abdomen (q = 3.67, p = 0.0369), but not thorax + head, surface areas (q = 3.21, p = 0.09), compared to females.

**Table 1.** Differences between male morphs and females in their surface area and body mass (n = 10; One-way ANOVAs with Tukey’s MCT). Different letters indicate statistically significant differences (p < 0.05), among morphs/sexes in that variable.

| Variable | Morph/Sex | Mean ± SD | Range |
| --- | --- | --- | --- |
| *Abdomen –* |  |  |  |
| *Surface Area (mm<sup>2</sup>)* | LM<sup>a</sup> | 453.0 ± 34.9 | 399.9 – 510.3 |
| | SM<sup>b</sup> | 327.6 ± 27.0 | 292.9 – 368.6 |
| | F<sup>c</sup> | 368.1 ± 44.6 | 316.9 – 449.5 |
| *Thorax and Head –* |  |  |  |
| *Surface Area (mm<sup>2</sup>)* | LM<sup>a</sup> | 489.1 ± 45.3 | 421.8 – 584.1 |
| | SM<sup>b</sup> | 332.8 ± 29.4 | 271.1 – 360.7 |
| | F<sup>b</sup> | 370.9 ± 40.4 | 302.5 – 431.6 |
| *Wet Body Mass (g)* |  |  |  |
| | LM<sup>a</sup> | 0.27 ± 0.05 | 0.19 – 0.34 |
| | SM<sup>c</sup> | 0.14 ± 0.02 | 0.11 – 0.19 |
| | F<sup>b</sup> | 0.19 ± 0.03 | 0.14 – 0.23 |

Hair density also differed across the abdomen and thorax dorsal surfaces based on sex/morph (Table 2, Fig 3; One-way ANOVA, abdomen: F = 59.14, df = 27, p < 0.0001; Kruskal-Wallis, thorax: K-W = 11.44, p = 0.0033). Large-morph males had more densely hairy abdomens and thoraxes than small-morph males (thorax: Z = 3.23, p = 0.0037; abdomen: q = 3.84, df = 27, p = 0.0299). Females did not differ from small-morph males in thorax hair density (Z = 0.74, p > 0.99), but had higher abdomen hair density compared to both male morphs (large: q = 10.98, p < 0.0001; small: q = 14.82, p < 0.0001).

**Fig 3.**
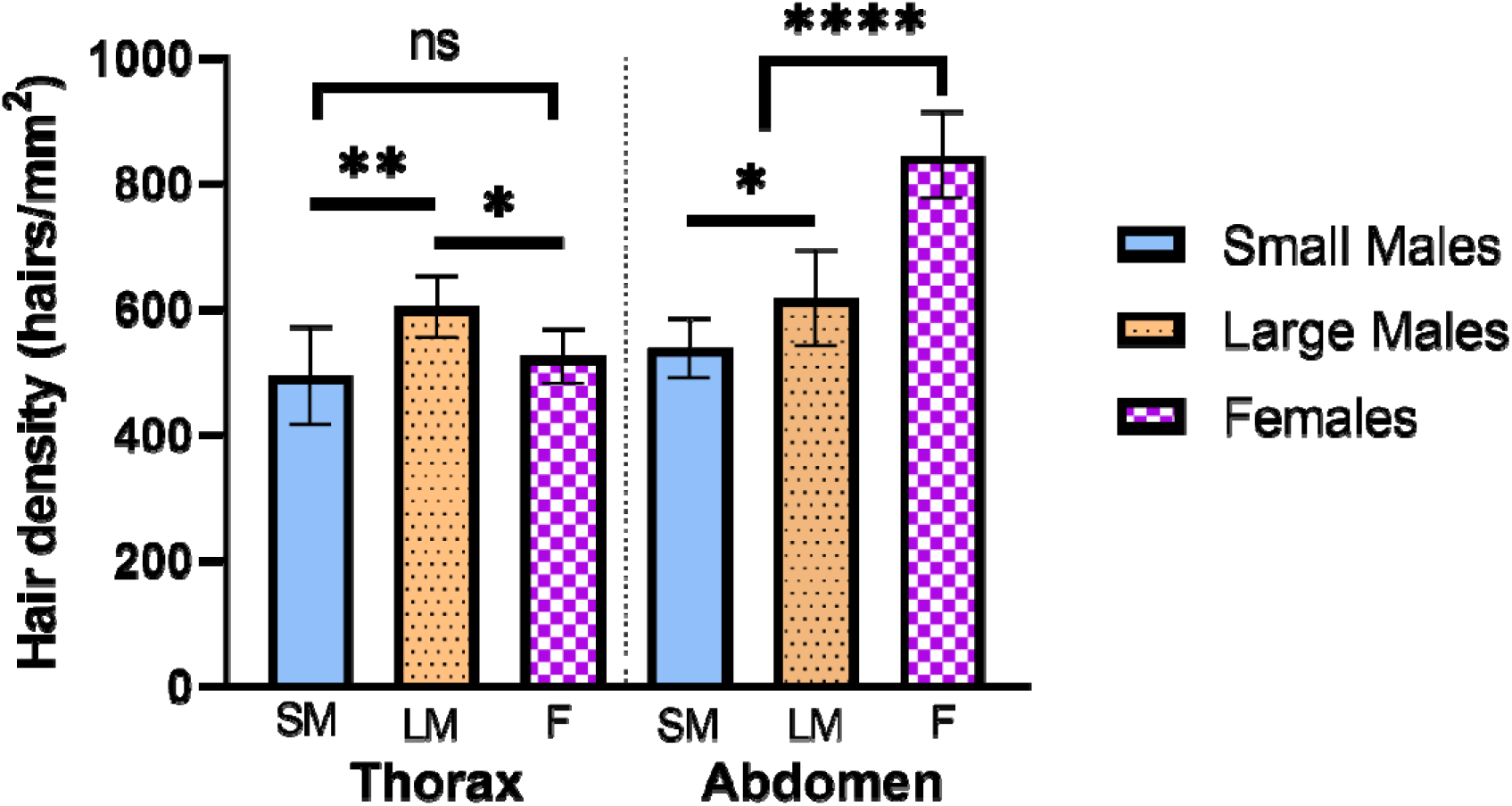
Differences in hair density of females, large and small morph males, on the dorsal surface. Large morph males have higher thorax hair density compared to small morph males and females (which do not differ in hair density; Kruskal-Wallis: K-W = 11.44, p = 0.0033). Females have higher abdominal hair density compared to males, and large-morph males have higher abdominal hair density compared to small-morph males (One-way ANOVA: F = 59.14, df = 27, p < 0.0001). Tukey’s or Dunn’s MCT: **** = p < 0.0001; ** p < 0.01; * = p < 0.05; ns = not significant

**Table 2.** Differences between male morphs and females in the density of hairs per mm^2^ on their dorsal thorax (Kruskal-Wallis with Dunn’s MCT) and abdominal surfaces (One-way ANOVA with Tukey’s MCT). Letters indicate significant differences (p < 0.05) across morphs/sex (n = 10) for that region. Asterisk indicates significant differences (p<0.05) between regions (abdomen vs. thorax) for that morph/sex (Kruskal-Wallis with Dunn’s MCT).

| Region | Morph/Sex | Mean ± SD | Range |
| --- | --- | --- | --- |
| *Abdomen - Dorsal* |  |  |  |
| | LM<sup>a</sup> | $617.8 \pm 76.82$ | $522.9 - 776.0$ |
| | SM<sup>b</sup> | $538.2 \pm 46.90$ | $479.6 - 626.1$ |
| | F<sup>c*</sup> | $845.6 \pm 69.37$ | $766.0 - 955.8$ |
| *Thorax - Dorsal* |  |  |  |
| | LM<sup>a</sup> | $604.6 \pm 48.86$ | $482.0 - 661.1$ |
| | SM<sup>b</sup> | $494.4 \pm 78.47$ | $411.5 - 656.1$ |
| | F<sup>b*</sup> | $525.7 \pm 44.20$ | $471.2 - 604.0$ |

Females had higher hair density on their abdomens compared to their thoraxes (Kruskal-Wallis, K-W = 38.14, p < 0.0001; Dunn’s MCT: Z = 4.67, p < 0.0001). There was no regional variation (e.g., thorax v. abdomen) in hair density within small-morph males, or large-morph males (Dunn’s MCT: small: Z = 0.92, p > 0.99; large: Z = 0.12, p > 0.99).

### Reflectance of Solar Radiation

Morph/sex significantly affected mean reflectance on the dorsal abdomen and thorax surfaces (Fig 4, Table 3, S1 Table; Two-way RM ANOVA; dorsal-abdomen: F = 26.32, df = 2, p < 0.0001; dorsal-thorax-unshaved: F = 22.01, df = 2, p < 0.0001; dorsal-thorax-shaved: F = 3.94, df = 2, p = 0.0314). Large-morph males had higher mean reflectance across the entire UV-NIR spectrum compared to small-morph males on both the dorsal abdomen and thorax surfaces (Fig 4A,B, S2 Table; Tukey’s: all p > 0.05).

**Fig 4.**
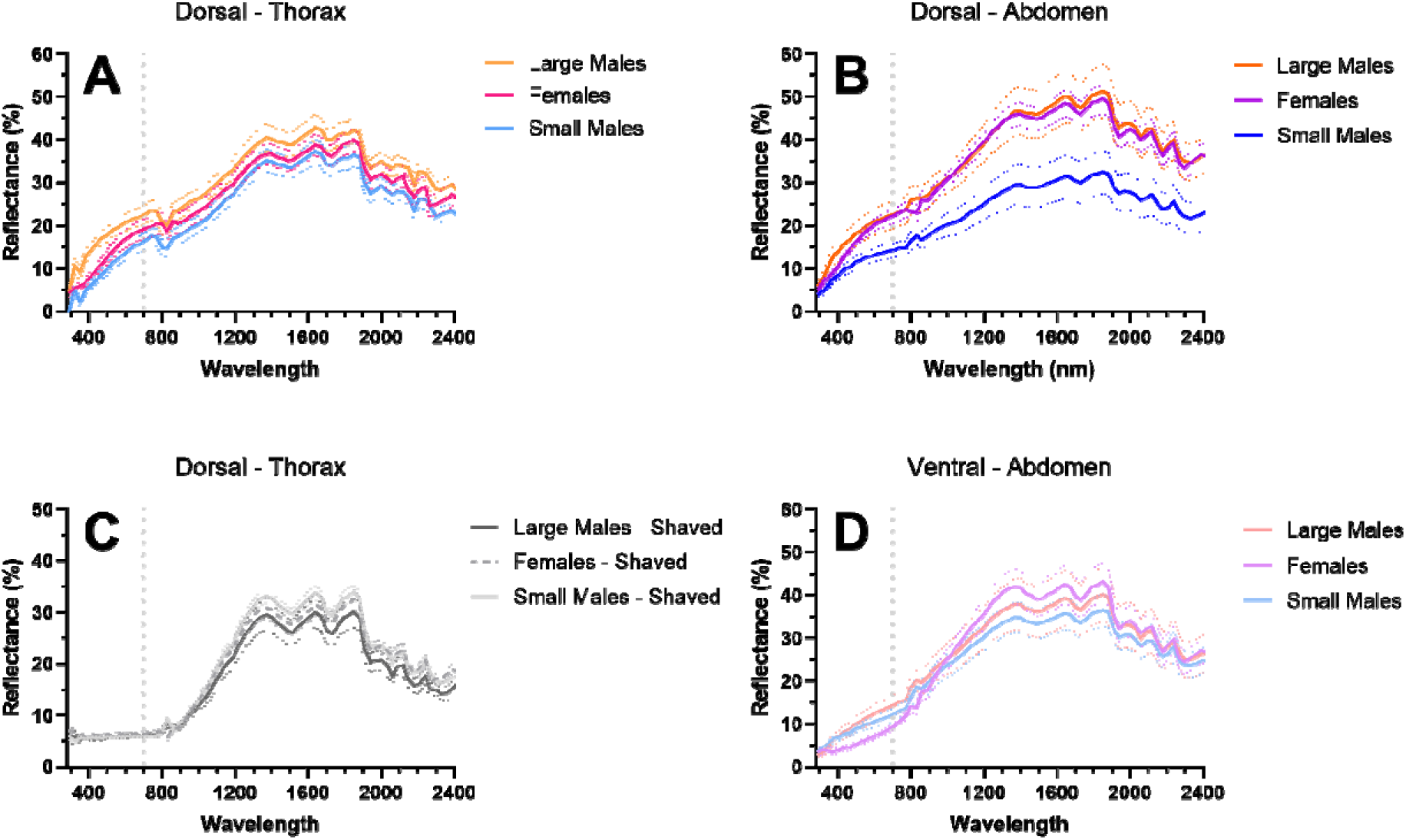
Reflectance of females, large males, and small males on the dorsal and ventral abdominal surface and dorsal thorax surface when shaved or unshaved. A) Differences in reflectance between large males, small males, and females on the dorsal surface of the thorax (Two-way RM ANOVA: F = 22.01, df = 2, p < 0.0001; MCT: S1 Table) and B) of the abdomen (F = 26.32, df = 2, p < 0.0001). C) The only significant difference in reflectance on the shaved dorsal thorax (e.g., cuticle-only) was between small males and females in the UV (F = 3.94, df = 2, p = 0.0314; MCT: UV SM-Female, q = 5.70, df = 16.66, p = 0.0301; all other comparisons, p > 0.05). D) There was no difference in mean reflectance between male morphs and females on the ventral surface of the abdomen (F = 0.65, df = 2, p = 0.53). Thick, solid or dashed lines represent mean reflectance; dotted lines are SD. The dotted, vertical line at 700 nm separates the VIS and NIR wavelengths.

**Table 3.**
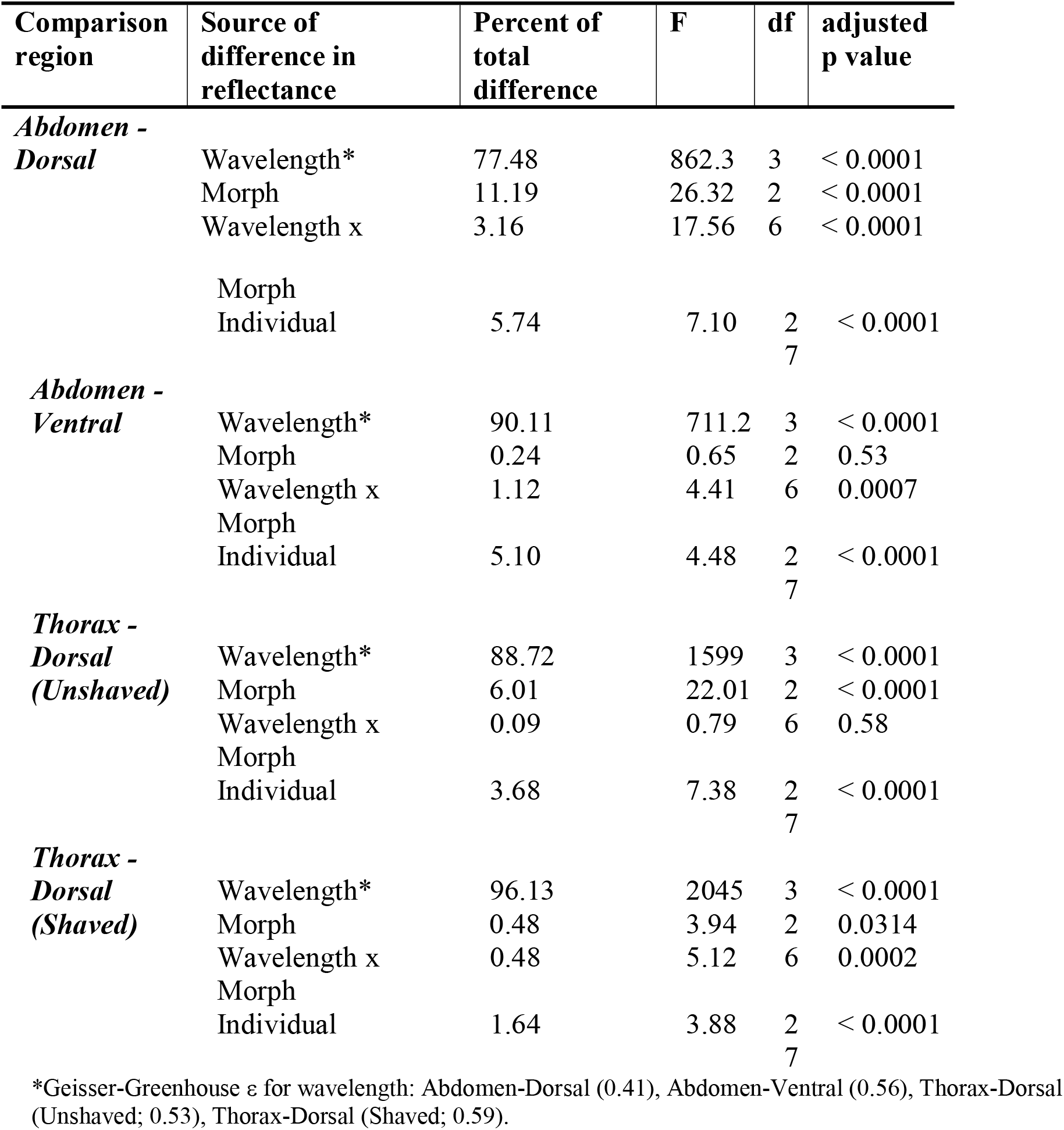
Two-way RM ANOVA with Geisser-Greenhouse Correction of mean reflectance of different morphs (small males, large males, and females) in different wavelength (UV [290-399], VIS [400-700], cNIR [701-1400] and fNIR [1401-2500]) on dorsal surface of the abdomen and thorax (unshaved), the dorsal surface of the thorax (shaved), and the ventral surface of the abdomen.

The only significant difference between morphs/sexes in the mean reflectance of the shaved dorsal thorax (e.g., cuticle-only) was between small males and females in the UV (Fig 4C, S2 Table; Tukey’s: q = 5.70, df = 16.66, p = 0.0301). All differences in mean reflectance between large and small males on the thorax disappeared when the hairs were shaved (S2 Table; Tukey’s: all p > 0.05). The presence of thorax hairs significantly increased mean reflectance across the entire UV-NIR spectrum for both small males and large males compared to just the cuticle (Table 4 & S3 Table; Two-way RM ANOVA with Sidak’s MCT: all p < 0.05).

**Table 4.**
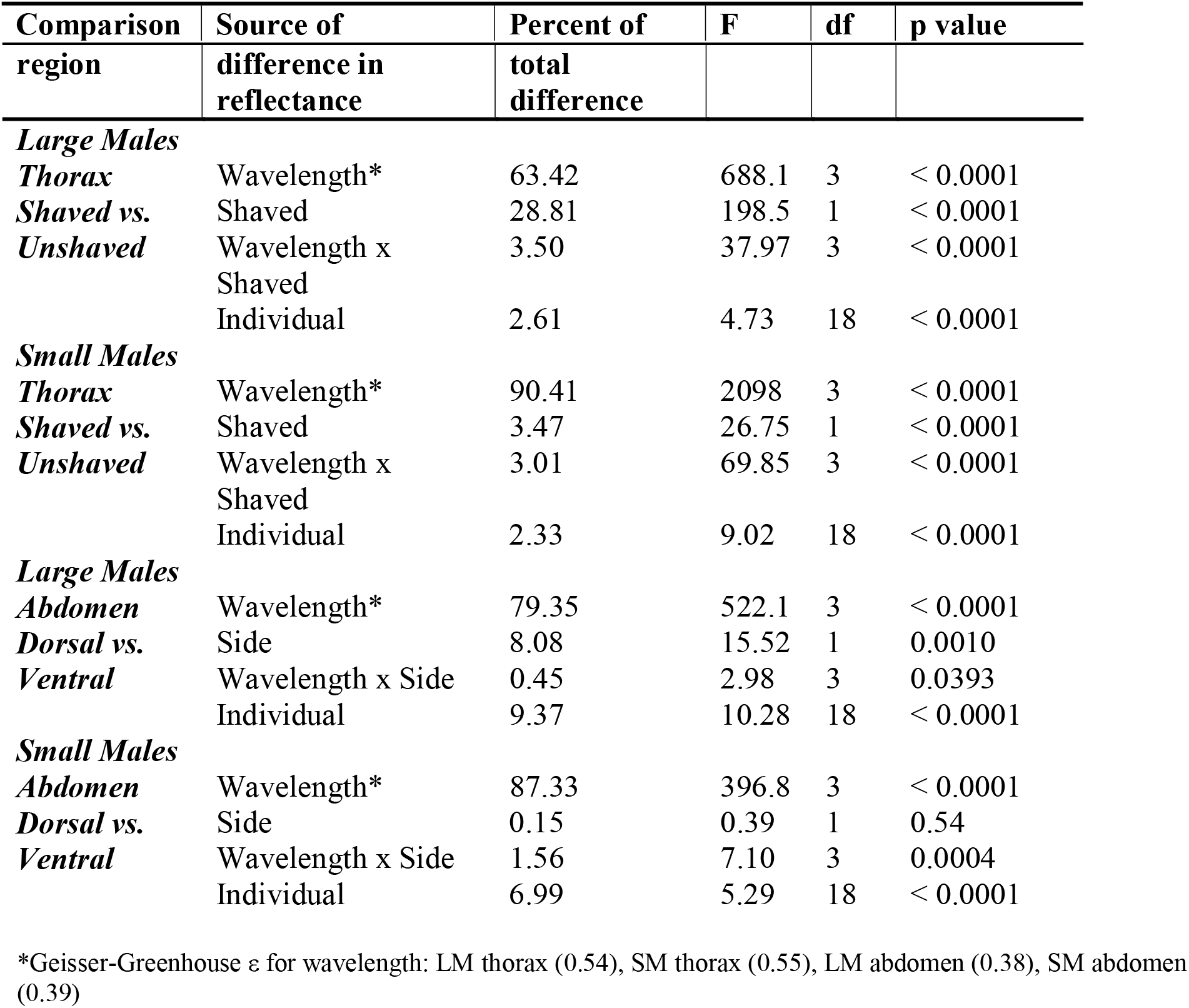
Two-way RM ANOVA with Geisser-Greenhouse Correction of mean reflectance of the thorax, shaved versus unshaved, and abdomen, dorsal versus ventral, at different wavelengths (UV [290-399], VIS [400-700], cNIR [701-1400] and fNIR [1401-2500]) in large males or small males.

The mean reflectance of the ventral abdominal surface did not differ across morphs/sexes (S2D Fig, Table 3; Two-way RM ANOVA: F = 0.65, df = 2, p = 0.53). The ventral surface of the abdomen of large males had lower mean reflectance across the entire UV-NIR spectrum compared to the dorsal surface (Table 4 & S3 Table; Two-way RM ANOVA with Sidak’s MCT: all p < 0.05), however the mean reflectance of the dorsal and ventral surface did not differ in small males (F = 0.39, df = 1, p = 0.54).

Sex/morph was dropped from the best fit model for both mean cNIR and fNIR reflectance on the unshaved, dorsal surface of the thorax after accounting for a correlation with VIS reflectance (ANOVA, cNIR: F = 1.00, p = 0.38; fNIR: F = 1.13, p = 0.34). Mean VIS and cNIR/fNIR reflectance were found to be highly correlated on the thorax surface (linear model, cNIR: F = 76.27, R^2^ = 0.72, p < 0.0001; fNIR: F = 44.6, R^2^ = 0.60, p < 0.0001). However, on the abdomen surface, sex/morph was a significant predictor variable for both mean cNIR (ANOVA, F = 6.28, p = 0.0059) and fNIR (F = 3.37, p = 0.0498) reflectance after accounting for a strong correlation with mean VIS reflectance (Table 5). However, large males and small males did not differ in abdomen surface reflectance after correcting for correlation with VIS reflectance (Tukey’s MCT; cNIR: p = 0.53; fNIR: p = 0.51).

**Table 5.** Linear models of the effects of sex/morph and mean VIS reflectance on cNIR [701-1400] and fNIR [1401-2500] reflectance on the dorsal surface of the abdomen.

| Comparison region | Effect | F | p value |
| --- | --- | --- | --- |
| cNIR | Mean VIS reflectance | 169.99 | < 0.0001 |
|  | Sex/morph | 6.28 | 0.0059 |
| fNIR | Mean VIS reflectance | 82.16 | < 0.0001 |
|  | Sex/morph | 3.37 | 0.0498 |

### Absorption of Radiation

The direct beam absorption coefficients of the dorsal thorax and abdominal surfaces of large-morph males were 0.76 ± 0.02 and 0.76 ± 0.06, significantly lower than those of small-morph males, which were 0.83 ± 0.02 and 0.82 ± 0.04 (Fig 5A and 6A, B; One-way ANOVAs, Tukey MCT, Thorax: q = 8.50, df = 27, p < 0.0001; Abdomen: q = 3.70, df = 27, p = 0.0372). Females had an abdominal absorption coefficient of 0.75 ± 0.02, and a thorax absorption coefficient of 0.80 ± 0.03. Their abdominal absorption coefficient was statistically similar to large males (One-way ANOVA, Tukey MCT: q = 0.50, df = 27, p = 0.93), and their thorax absorption coefficient was higher than large males (q = 4.91, df = 27, p = 0.0048). Their abdominal (q = 4.20, df = 27, p = 0.0165) and thorax (q = 3.58, df = 27, p = 0.0444) absorption coefficients were lower than small males (though differences in thorax coefficients were marginally statistically significant).

**Fig 5.**
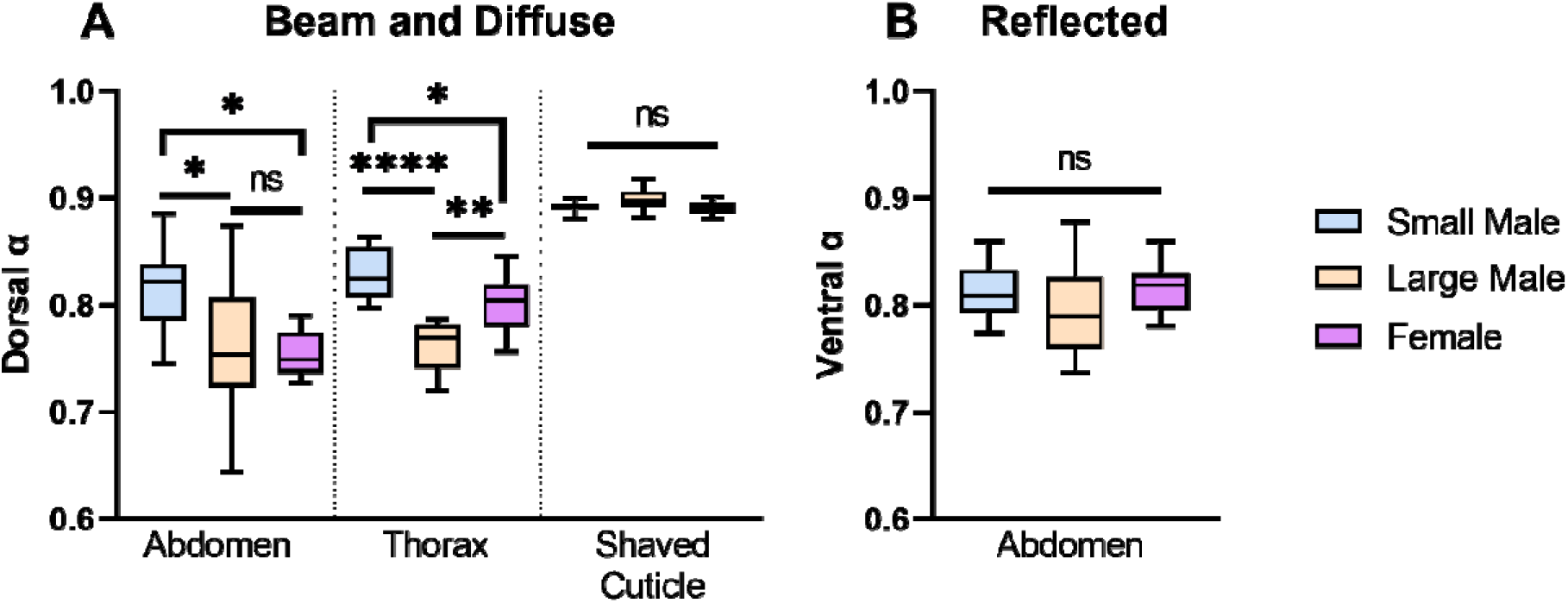
Difference in direct beam and diffuse absorption coefficients of the dorsal abdomen and thorax between male morphs. A) Difference in absorption of direct beam and diffuse sky solar radiation of the dorsal abdomen, thorax, and shaved thorax cuticular surfaces. Large males and females have lower absorption coefficients compared to small-morph males on the thorax and abdomen when hairs are present, but when hairs are shaved the absorption coefficients do not vary across male morphs and females. B) There is no difference in the absorption coefficients of male morphs and females when considering reflected solar radiation off a light sand substrate on the ventral surface. **** = p < 0.0001; ** p < 0.01; * = p < 0.05; ns = not significant

**Fig 6.**
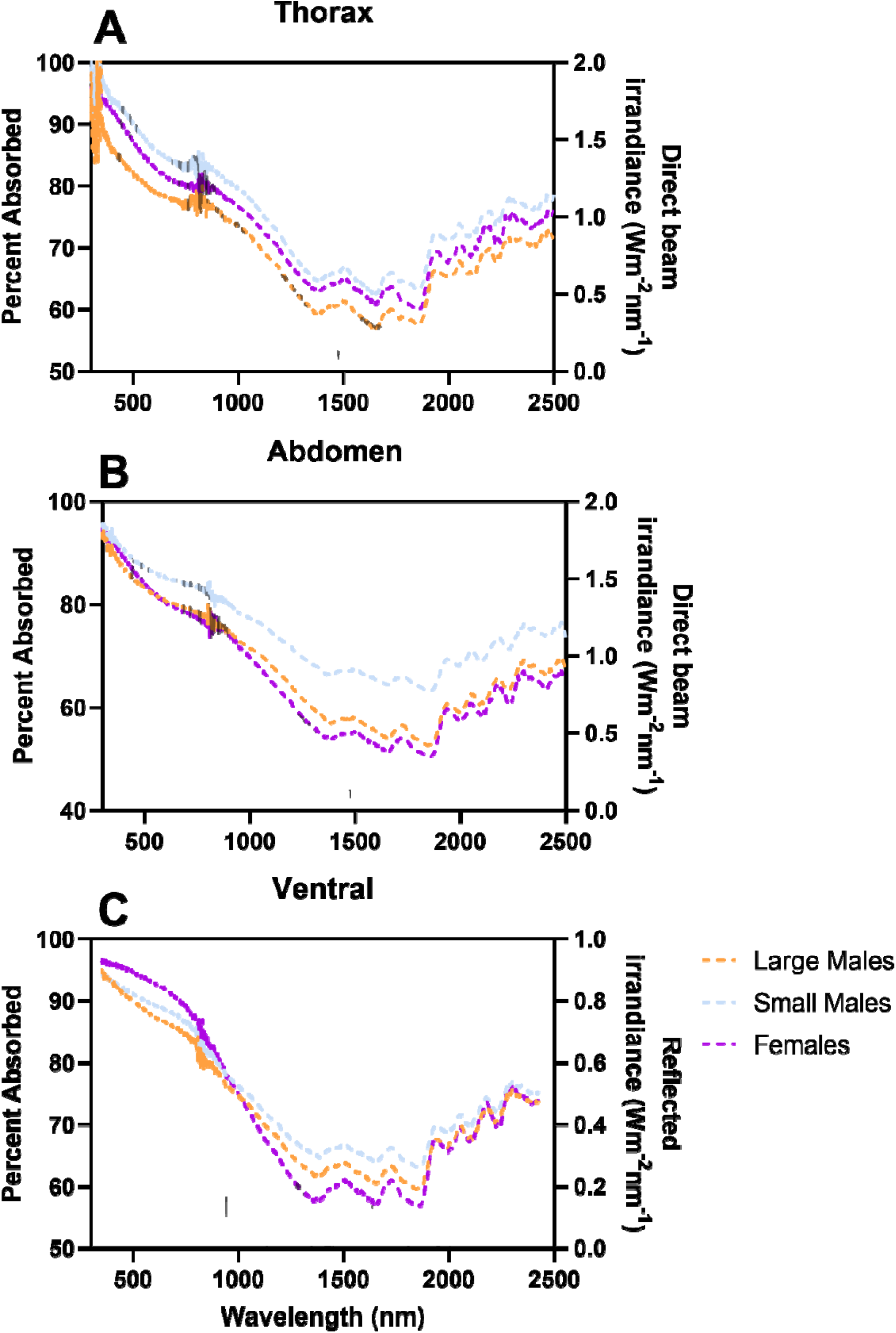
Spectral properties of *C. pallida* thorax and abdomen in relation to the irradiance spectrum of direct beam sunlight and sunlight reflected off light sand. Left y-axis represents the percentage of absorbed light at each wavelength, absorbed by the dorsal thorax (A), dorsal abdomen (B), or ventral abdomen (C) cuticle and hairs of *C. pallida* large- and small-males or females (mean, n = 10 each; females in purple, large males in orange, small males in blue). Right axis represents the solar beam irradiance based on solar irradiance distribution standard (Gueymard et al. 2002) in A and B, or the reflected irradiance off light sand (Kokaly et al. 2017) in C (all irradiance in grey).

Females and males did not differ in the direct beam absorption coefficients of their shaved cuticles (Fig 5A; One-way ANOVA, F = 2.78, df = 27, p = 0.08). There was also no difference in the absorption of reflected light by the ventral surfaces of male morphs and females (Fig 5B and 6C; One-way ANOVA, F = 1.21, df = 27, p = 0.31).

Direct beam and diffuse sky radiation was calculated to be 1,178.82 W/m^2^ at 1000 h on April 20^th^. Reflected radiative energy was higher when accounting for wavelength specific effects (329.85 W/m^2^) as compared to when applying the same albedo for light sand (0.245) across all wavelengths (288.81 W/m^2^); we used 329.85 W/m^2^ for all further calculations.

Large-morph males absorbed a mean of 0.36 ± 0.02 W of thermal energy from reflected, direct, and diffuse radiative sources (1.28 ± 0.13 W/g of body mass), compared to 0.26 ± 0.02 W for small-morph males (1.85 ± 0.21 W/g of body mass). If large-morph males had the same reflectance profiles as the averages for small-morph males, they would absorb 0.37 ± 0.03 W of thermal energy (or 1.40 ± 0.15 W/g of body mass, an 8-9% increase in W/g).

Females absorbed a mean 0.28 ± 0.03 W of thermal energy, or 1.40 ± 0.11 W/g of body mass. For all three groups, hairs on the dorsal surface reduced thermal heat flux due to solar radiation; thermal flux for shaved large males was 0.41 ± 0.03 W (1.42 ± 0.17 W/g), for small males was 0.28 ± 0.02 W (1.94 ± 0.23 W/g), and for females was 0.31 ± 0.03 W (1.63 ± 0.12 W/g).

## Discussion

Comparative studies of animal solar reflectance (or VIS coloration) often collect data at a macrogeographic scale, and demonstrate the effect of geographic gradients in climate variables (such as ambient temperature or solar radiation) on solar reflectance [2,4,11,16,19]. However, large differences in ambient temperature can occur within populations when individuals use different microclimates. The 1 cm above-ground microclimate where large male *C. pallida* engaged in their mating behaviors has air temperatures > 8 °C hotter than the 1 m above-ground hovering microclimate used by small males by 1100 am [32], demonstrating that there can be substantial differences in thermal selective pressure across distinct microclimates at the same site. The reduced solar absorption of large-morph males in both the VIS and NIR may be a unique thermal adaptation to this very hot microclimate, where they must behave for prolonged periods to successfully access mates (males are known to dig and fight for up to 19 minutes [28]).

Morph-specific variation in solar reflectance on the dorsal surface went beyond the VIS, with consistent differences in reflectance in the NIR. Because morph reflectance differences were strongly correlated in the VIS and NIR, selection on visual characteristics (e.g., anti-predator adaptations) could indirectly affect NIR reflectance. However, the higher UV-NIR reflectance of large-morph males conferred a thermal benefit for this morph, which utilizes a hotter microclimate: the higher UV-NIR reflectance of large-morph males led to an 8-9% reduction in W/g of absorbed solar energy. We suggest that morph differences in solar reflectance may be, in part, a thermoregulatory adaptation to their distinct microclimates; reflectance differences beyond visible coloration suggests there may be non-visual selective factors partially or wholly driving reflectance differences.

Females had intermediate dorsal absorption coefficients compared to males: female thorax reflectance profiles were similar to small males, but lower compared to large males, and abdomen reflectance profiles were similar to large males, but higher compared to small males. There was no significant difference in the absorption of reflected irradiance on the ventral surface among morphs and sexes, supporting the hypothesis that differences in reflectance may be an adaptation that functions to increase radiative heat gain in cooler microclimates, and decrease it in hotter microclimates.

Differences in dorsal reflectance were entirely caused by the hairs; there was no significant difference in the shaved cuticular absorption coefficients of the male morphs or females. Large-morph males had higher hair densities on the thorax and abdominal surfaces compared to small-morph males; however, they had decreased hair density on the abdomen compared to females, despite similar reflectance profiles. This suggests that both hair density, as well as structural features of the hairs (S2 Fig), may contribute to the reflectance profile differences across male morphs and females. Microstructural features of insect hairs and scales are known to influence their reflective properties in both the NIR and mid-infrared [6,8,46]. The striations on the external surface of the hair (S2 Fig) are reminiscent of the hairs of the Saharan silver ant [8]. The striated external surface structure of the ant’s hairs contributes to total internal reflection; however, the lack of triangular shape, and the orientation of the hairs as perpendicular (as compared to parallel) to the cuticle, reduce the likelihood of this structural mechanism in the *C. pallida* case. Alternately, the hollow structure of the hair may play a role in thermal management, as is the case in polar bear fur [47] and the silver ant [8].

Temperature can vary both spatially and temporally at a given site: across the course of the morning, temperature increased from a low of 16.9 °C to a high of 44.5 °C between 0600 and 1100 hrs in the 1 cm microclimate, and from 16.1 °C to 38.9 °C in the 1 m microclimate [32]. In male *Colias* butterflies, intraspecific increases in radiative heat gain due to melanization allowed for longer flight periods at cooler air temperatures, increasing access to mating opportunities [48–49]. The mean body size of the *C. pallida* male population increases throughout the morning (both for males foraging, and males engaged in mating-relevant behaviors [50]), suggesting smaller males may use their darker coloration to increase radiative heat gain and their access to mates during cooler early morning periods when large males are not yet as prevalent in the population.

An additional implication of this work relates to unraveling the various factors that may lead to stabilizing selection for dimorphic body morphologies in alternative reproductive tactic systems. Generally, it has been hypothesized that bird predation may select against the large-morph male mating advantage in the *C. pallida* system, producing persistent size variation and two distinct morphs [34]. However, bird predation is variable across field sites [34] while size dimorphism is not. In addition, large-morph males are not behaviorally wary of predation events (which might be expected if predation was as a significant selective force); for example, large-morph males can be captured easily with just the fingertips and will continue to dig or fight when experimenter shadows pass over them [31, Barrett pers. obs.]. An alternate hypothesis, better supported by our data, relates the continued existence of two microclimate-specialized morphs to the thermoregulatory selective pressures of the available microclimates. Males of an intermediate size and UV-NIR reflectance profile would be disadvantaged due to higher metabolic rates (and thus higher metabolic heat production) when using small male mating strategies (in other bees with intraspecific body size variation, smaller individuals have increased power efficiency in flight without any additional metabolic cost [51]), while also disadvantaged due to their increased solar absorptivity when using the large male mating strategies. Individual morphological specialization in relation to body form and reflectance may potentially facilitate the stability of the male *C. pallida* alternative reproductive tactic system.

Our study demonstrates the importance of both VIS and NIR reflectance as intraspecifically traits conferring a thermal benefit to large-morph males in the hotter microclimate. Variation in reflectance of shortwave radiation, and other morphological adaptation to that benefit organisms facing thermal pressures (such as larger body sizes), may alter the necessity for physiological thermoregulatory differences – for example, large morph males that can avoid overheating in the sun due to their reduced relative solar heat load may not need higher thermal tolerances to survive (see [32]). The interplay between these morphological and physiological thermoregulatory benefits/strategies thus deserve greater attention. In addition, this variation suggests individuals within populations may find their fitness-relevant behaviors to be differently constrained by increasing global temperatures due to the effects of variation in shortwave radiative heat gain on their energy budget. If thermoregulatory selective pressures play a role in maintaining size or behavioral variation within species (such as alternative reproductive tactic systems), this suggests that climate change may have impacts that stretch beyond species-level diversity – affecting intraspecific morphological and behavioral diversity as well.

We demonstrate that individual differences, often overlooked, may be important for understanding the selective pressures acting on individuals to generate evolutionary adaptations to fine-scale ecological variation. Future studies should thus pay greater attention to the effects of morphological, physiological, or behavioral variation between conspecifics, which may have important consequences for understanding proximate and ultimate causes of ecological reflectance and coloration patterns.

## Acknowledgements

This work would not have been possible without the support of Kathryn Busby, Logan Schoolcraft, and Stephen Buchmann. Offir Cohen assisted with making the aperture, and spectrophotometer training. MB was not a National Science Foundation postdoctoral fellow while completing this work but is an NSF postdoctoral fellow at the time of publication: any opinions, findings, conclusions, or recommendations expressed in this manuscript are the authors, and do not necessarily reflect the views of the NSF.

## Supporting information

**S1 Fig.**
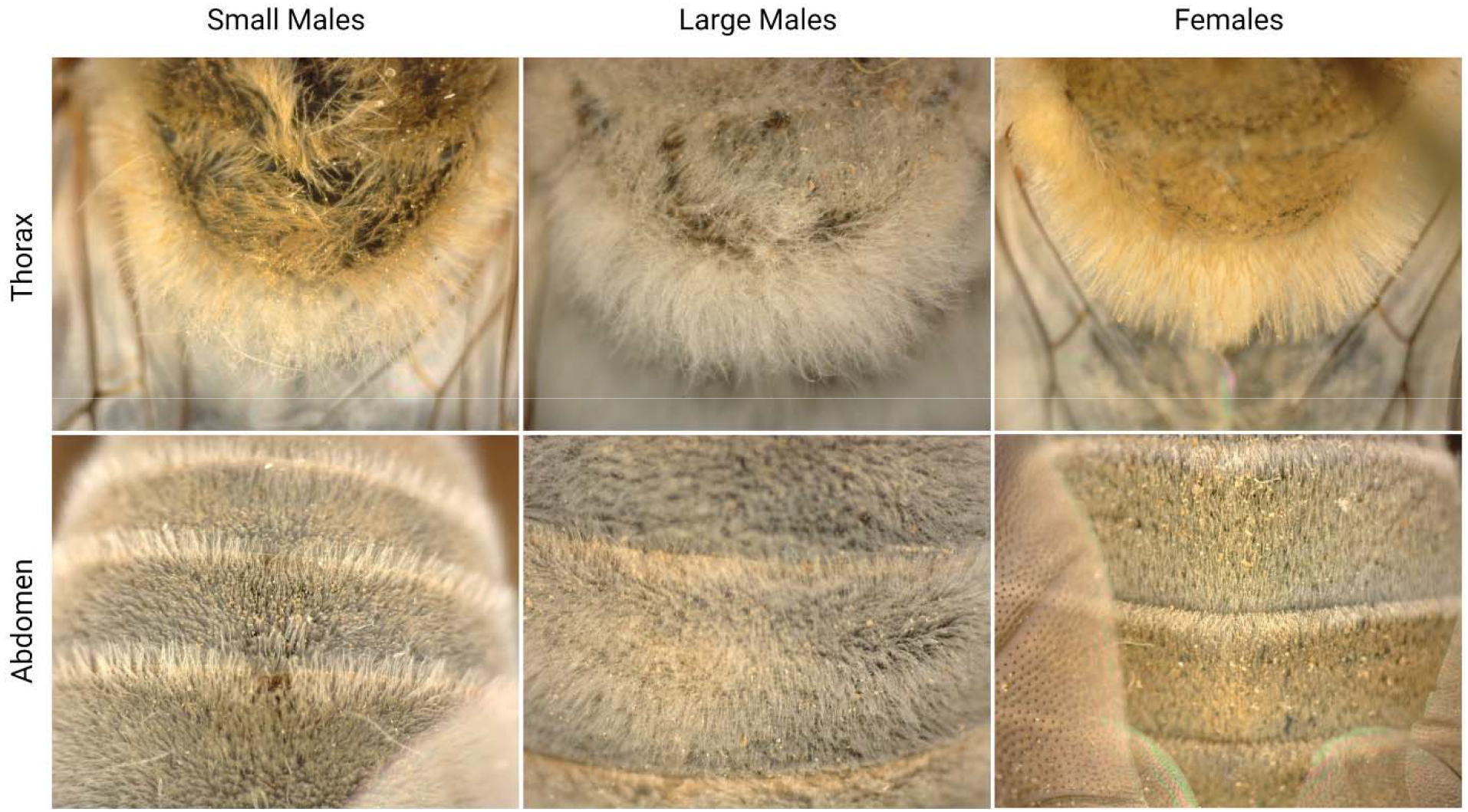
Difference in coloration of *C. pallida* hairs on the thoraxes and abdomens of large- and small-morph males and females. All photos taken at 85X magnification on a DinoLite AM4915ZT.

**S2 Fig.**
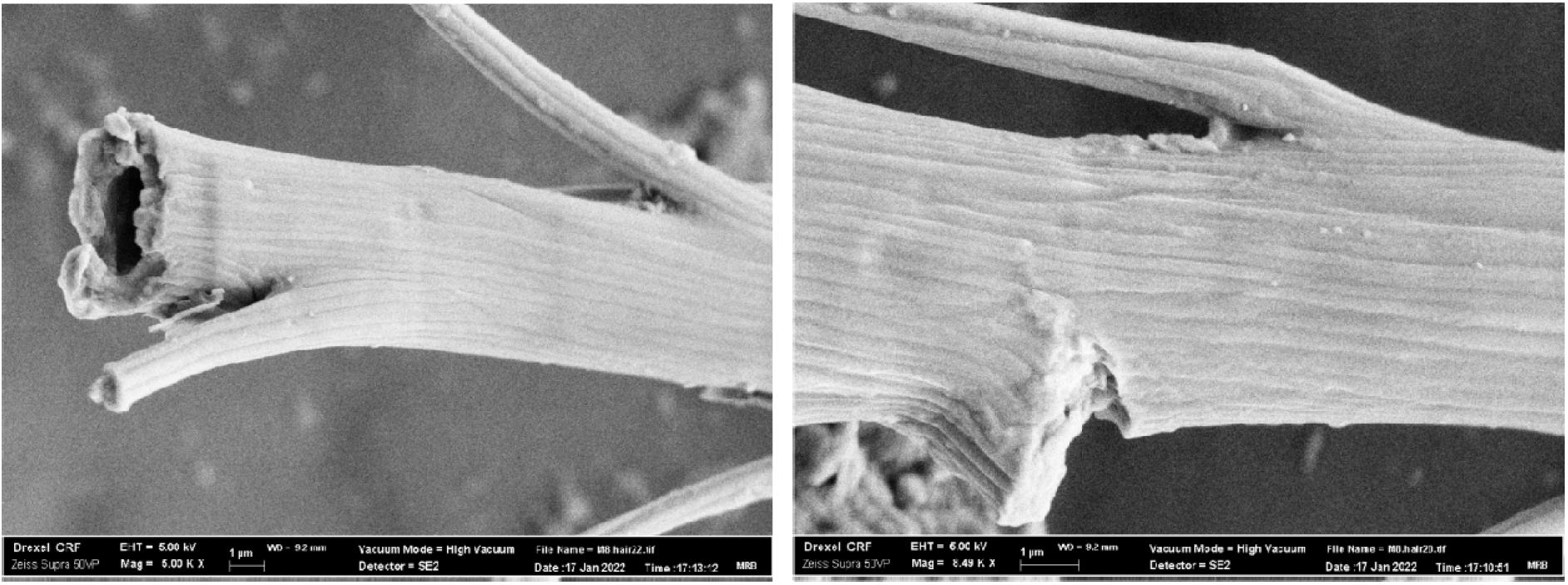
Large male thorax hair characteristics. **Left)** The hairs of large-morph males have a round cross-section, and are covered in striations (5000X). **Right)** Close-up of striations on the exterior surface of large-morph male thorax hairs (8500X). The somewhat regular striations are reminiscent of the structure of the highly reflective hairs of the Saharan silver ant (in Shi et al. 2015).

**S1 Table.**
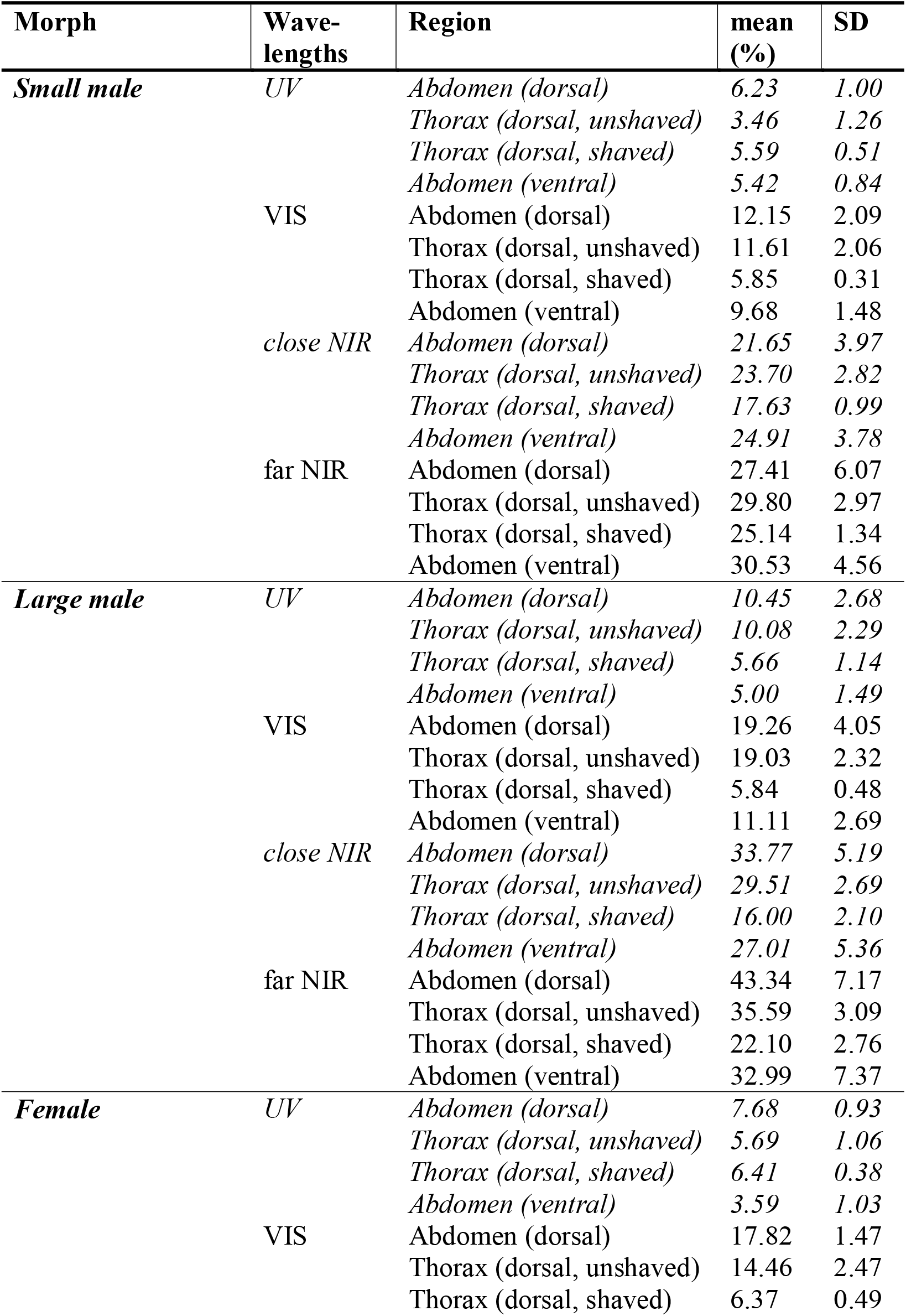
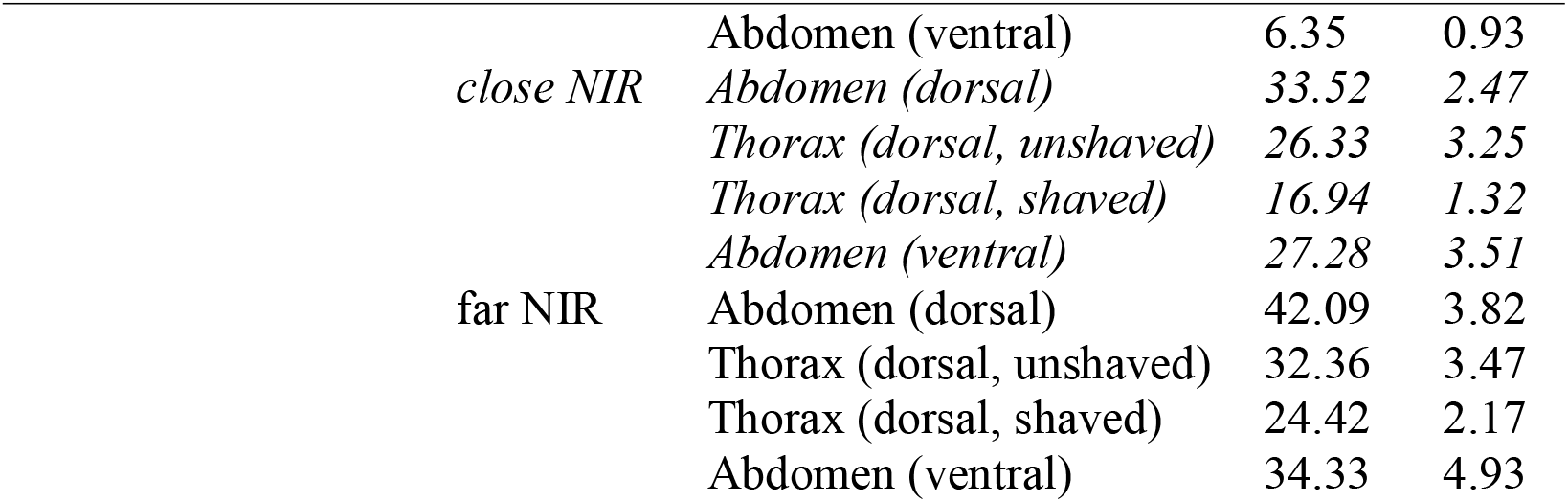
Mean percent reflectance of males and females in the UV-NIR. Mean ± SD of mean percent reflectance of small males, large males, and females in the UV (290-399), VIS (400-700), close NIR (701-1400) and far NIR (1401-2500) on dorsal surface of the abdomen and thorax (unshaved and shaved).

**S2 Table.**
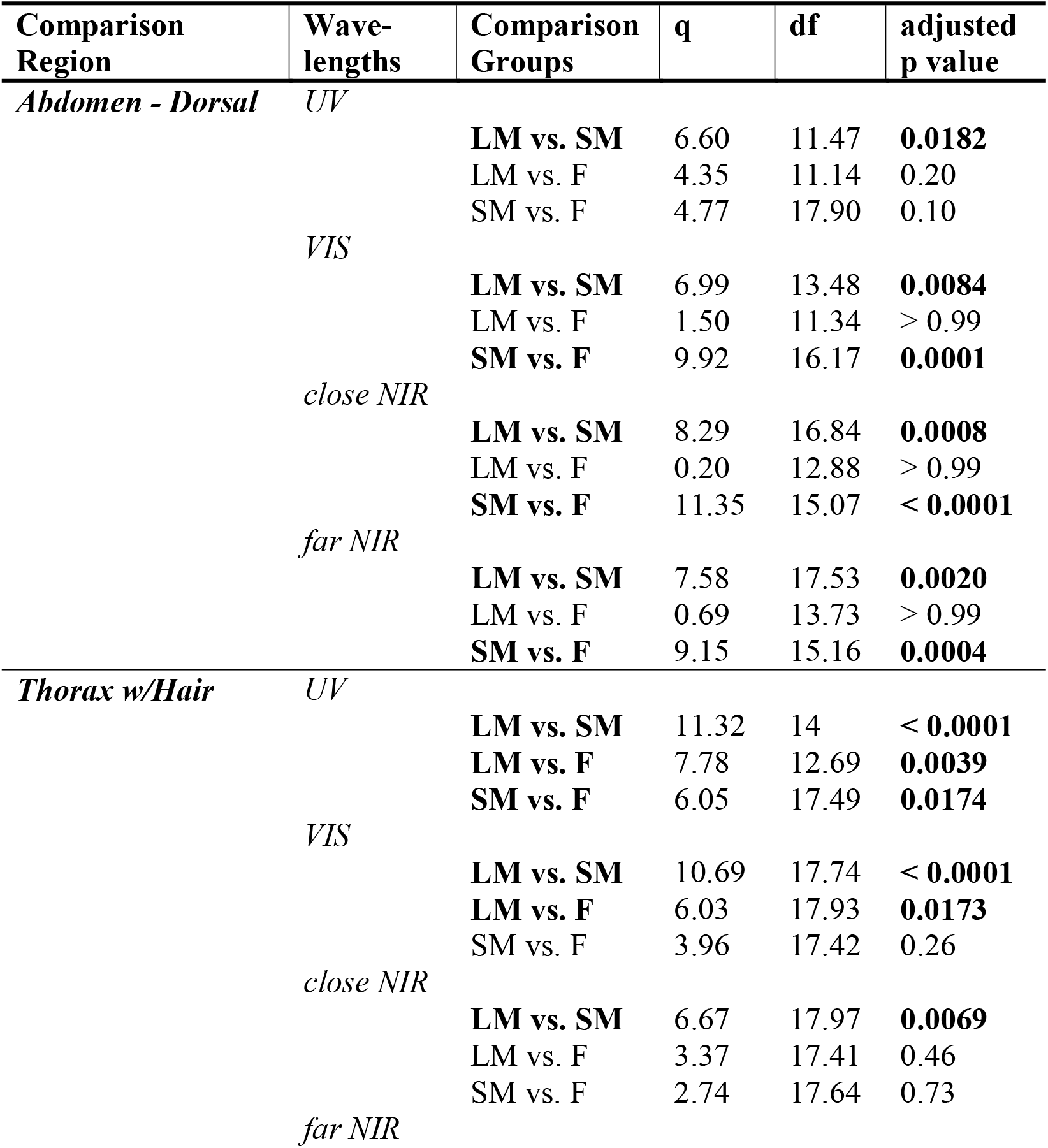
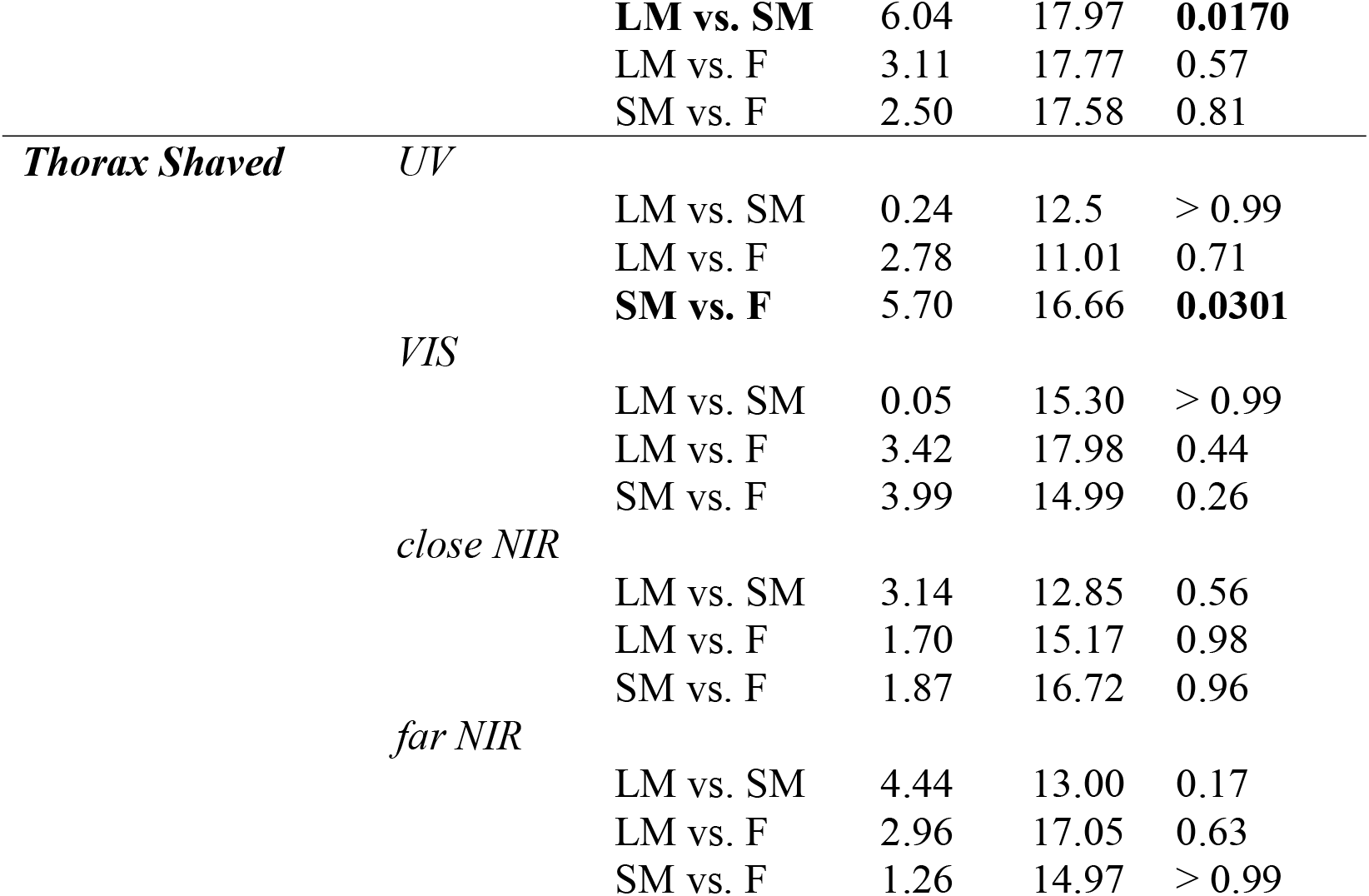
Tukey’s multiple comparisons test (following two-way RM ANOVA with Geisser-Greenhouse Correction) of mean reflectance of small males (SM), large males (LM), and females (F) in the UV (290-399), VIS (400-700), close NIR (701-1400) and far NIR (1401-2500) on dorsal surface of the abdomen and thorax (unshaved and shaved).

**S3 Table.** Sidak’s multiple comparisons test (following two-way RM ANOVA with Geisser-Greenhouse Correction) of mean reflectance of the thorax, shaved versus unshaved, and abdomen, dorsal versus ventral, at different wavelengths (UV [290-399], VIS [400-700], close NIR [701-1400] and far NIR [1401-2500]) in large males or small males.

| **Morph** | **Comparison** | **Wave length** | **t** | **df** | **adjusted p value** |
| --- | --- | --- | --- | --- | --- |
| **Large Males** | Shaved vs. Unshaved |  |  |  |  |
| - |  |  |  |  |  |
| **Thorax** |  | UV | 5.46 | 13.22 | 0.0004 |
|  |  | VIS | 17.59 | 9.76 | < 0.0001 |
|  |  | cNIR | 12.52 | 16.97 | < 0.0001 |
|  |  | fNIR | 10.30 | 17.77 | < 0.0001 |
| **Small Males -** | Shaved vs. Unshaved |  |  |  |  |
| **Thorax** |  | UV | 4.95 | 11.91 | 0.0014 |
|  |  | VIS | 8.77 | 9.40 | < 0.0001 |
|  |  | cNIR | 6.43 | 11.21 | 0.0002 |
|  |  | fNIR | 4.53 | 12.50 | 0.0025 |
| **Large Males** | Dorsal vs. Ventral |  |  |  |  |
| - |  |  |  |  |  |
| **Abdomen** |  | UV | 5.61 | 14.06 | 0.0003 |
|  |  | VIS | 5.31 | 15.66 | 0.0003 |
|  |  | cNIR | 2.82 | 17.93 | 0.0450 |
|  |  | fNIR | 3.19 | 17.99 | 0.0204 |

## Notes

**Declarations of Interest:** none.

https://doi.org/10.5061/dryad.73n5tb31q

### Competing Interest Statement

The authors have declared no competing interest.

### Summary of Updates

Some revisions to the statistics, demonstrating high correlation between NIR and VIS reflectance. Figures updated for clarity and some supplementary files added to the main text. Added permit information.

